# Development of inducible promoter and CRISPRi plasmids functional in *Rickettsia rickettsii*

**DOI:** 10.1101/2024.06.12.598709

**Authors:** Adam M. Nock, Tina R. Clark, Ted Hackstadt

## Abstract

*Rickettsia rickettsii* is an obligate intracellular, tick-borne bacterium that causes Rocky Mountain spotted fever. The demanding nature of cultivating these bacteria within host cells and the labor involved in obtaining clonal isolates has severely limited progress regarding the development of compatible genetic tools to study this pathogen. Conditional expression of genes which might be toxic or have an otherwise undesirable effect is the next logical goal to expand upon the constitutive expression plasmids generated thus far. We describe the construction of an inducible promoter system based on the tet-On system, leveraging design elements from the anhydrotetracycline inducible promoter system used for *Borrelia burgdorferi* and one of the few characterized rickettsial promoters for the outer membrane gene, *rompB* (*sca5*). The functionality of this promoter is demonstrated via fluorescence of induced mScarlet production and was then used to construct a generalized inducible expression vector for *R. rickettsii*. The development of a functional inducible promoter was then applied to the construction of a CRISPR interference plasmid as a means to reduce or essentially silence the transcription of targeted genes. We demonstrate the viability of a simplified, single vector CRISPRi system to disrupt gene expression in *R. rickettsii* targeting the type IV secreted effector *rarP2* and autotransporter peptidase *rapL* as examples.

**IMPORTANCE:** This work expands upon the genetic toolbox available for *R. rickettsii*. This is the first report of both an inducible promoter and CRISPRi system compatible with *Rickettsia*, which may provide key instruments for the development of further tools. The development of an inducible promoter system allows for the overexpression of genes which might be toxic when expressed constitutively. The CRISPRi system enables the ability to knockdown genes with specificity, and critically, genes which may be essential and could not otherwise be knocked out. These developments may provide the foundation for unlocking genetic tools for other pathogens of the order Rickettsiales, such as the *Anaplasma*, *Orientia*, and *Ehrlichia* for which there are currently no inducible promoters or CRISPRi platforms.

## INTRODUCTION

*Rickettsia rickettsii* is a tick-borne pathogen causing Rocky Mountain spotted fever, a disease with a high mortality rate when untreated. Because *R. rickettsii* is obligately intracellular, and there is currently no axenic media available, progress on genetic tools has been slow compared to model organisms (1, 2). While the development of genetic tools for other intracellular pathogens such as *Chlamydia*, *Coxiella*, and *Mycobacteria* have accelerated, but progress in the rickettsiales has lagged. Key steps in the development of genetic tools for *Rickettsia* have included transformation and the implementation of fluorescent proteins, which allowed for transposon mutagenesis and selection (3, 4). Another critical leap forward occurred as rickettsial plasmids were discovered, described, and used to generate shuttle vectors greatly facilitating molecular cloning and subsequent transformation of rickettsia (5, 6). These tools have been widely adopted and adapted by the rickettsiology field to develop plasmids for constitutive overexpression and genetic knockout systems (7–9). The challenge which has remained and areas of improvement that require addressing are the reliability of these knockout systems and the challenges involved in dealing with genes of interest that are essential and cannot be knocked out, and finally overexpressing genes which may be toxic when overexpressed.

A critical element required to solve these issues is the development of an inducible system as has been achieved for other intracellular bacteria. The specific architecture of promoters and consequent dynamics of transcription is understudied in the rickettsiales compared to model systems such as *Escherichia coli* or even other alphaproteobacteria. Solving the promoter architecture and an understanding of transcriptional initiation and termination represent core hurdles to improved genetic tool development. However, some *Rickettsia* promoters have been described including those for citrate synthase (*gltA*), *rpsL*, *rompA* (*sca0*), and *rompB* (*sca5*) (10–12). Some transcriptional terminators in *R. prowazekii* have also been studied (13). The information gleaned from these studies has been and continues to be an important starting point for the construction and design of genetic tools for rickettsia.

Herein we describe the construction of an inducible-promoter based on the tet-On system compatible with *R. rickettsia* based on previously described promoters and rickettsial shuttle vectors to generate generalized vectors to achieve anhydrotetracycline-inducible overexpression of genes of interest (14). We then utilized this system to generate a Single plasmid vector for CRISPR interference (CRISPRi) leveraging dCas9, a catalytically dead version of Cas9, to knockdown the expression of target genes (15). CRISPRi systems leveraging inducible promoters have been developed for other intracellular pathogens including *Chlamydia trachomatis*, *Coxiella burnetii*, and *Mycobacteria tuberculosis* (16–18).

## RESULTS

### Generation of an inducible expression system compatible with *R. rickettsii*

Because using plasmids constitutively expressing rickettsial genes present occasional technical issues due to presumed toxicity of the genes expressed, we sought to generate a functional, rickettsia-compatible inducible expression system. An anhydrotetracycline-inducible system was opted for due to its success in other intracellular pathogens (19–21). The tolerance of *R. rickettsii* Sheila Smith in Vero cells to different concentrations of anhydrotetracycline were tested and concentrations up to 250 ng/mL were found to be tolerated (Fig 1A). A test-bed plasmid, pRAMtetS, based on shuttle vectors designed for *Rickettsia* (6) was generated with constitutively expressed mNeongreen (22) and mScarlet (23) under the control of a novel promoter including a tet operator (tetO) site (Fig 1B). The promoter, P_rrtetO_, is a combination of a portion of the *rompB* promoter, defined previously (11), and the original anhydrotetracycline inducible promoter designed for *Borrelia burgdorferi* (24), another tick-borne bacterium with low GC content (Fig 1C). The addition of anhydrotetracycline at 10 ng/mL to media was sufficient to induce expression of mScarlet by *R. rickettsii* harboring pRAMtetS infecting Vero cells compared to a DMSO vehicle control (Fig 1D). To build an inducible expression plasmid suitable for general use, pRAMin, we replaced the mScarlet gene with an NheI site for linearization such that genes of interest can be cloned into the expression site using NEBuilder HiFi DNA Assembly or Gibson assembly (Fig 1E) (25). A schematic of the development and intermediate plasmid constructs is depicted in Fig. S1.

**Figure 1:**
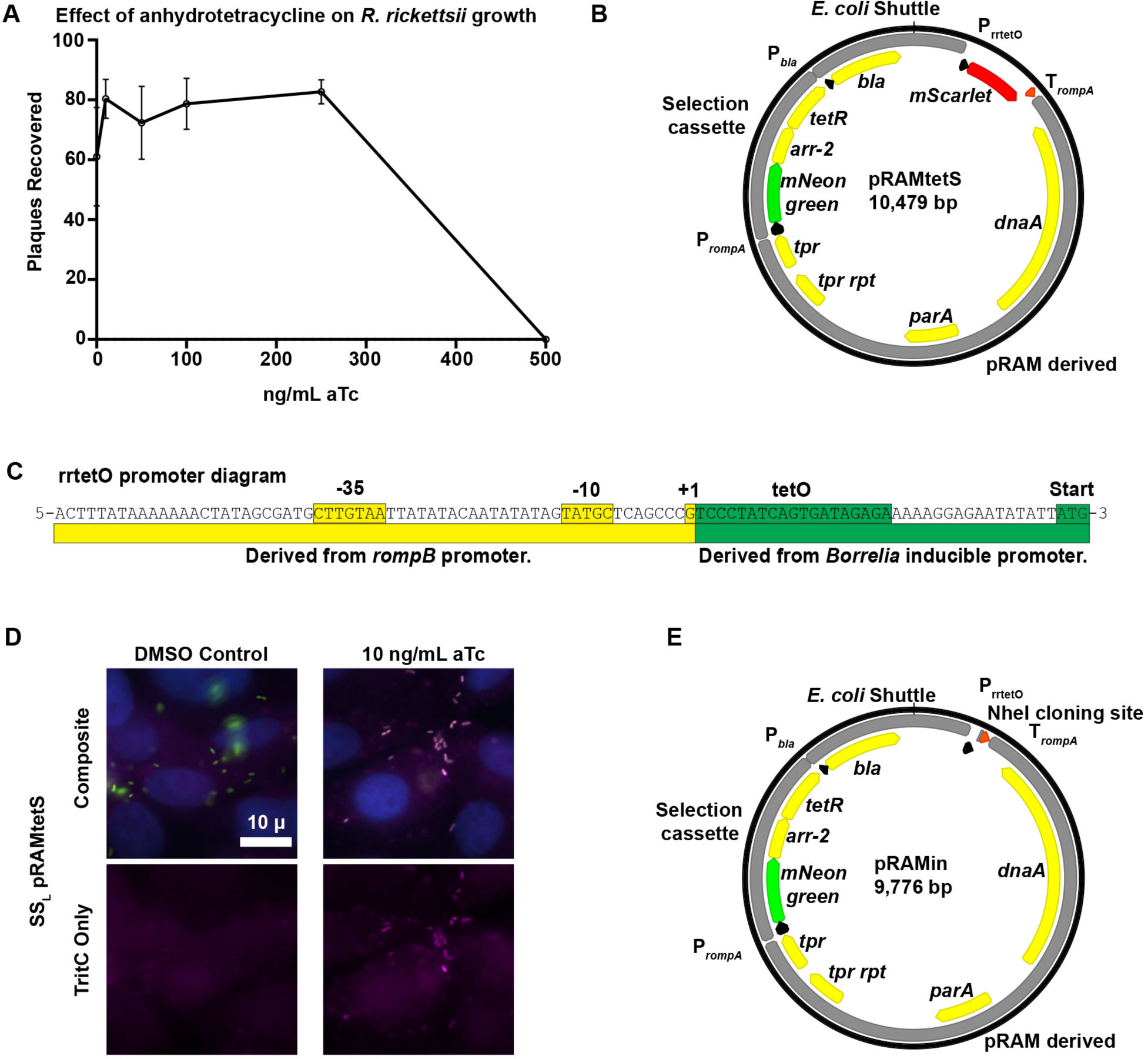
Development of an inducible promoter system compatible with *R. rickettsii*. (A) Effect of anhydrotetracycline on replication of *R. rickettsii*. (B) Plasmid map of pRAMtetS, used to test the functionality of P_rrtetO_ via aTc induction of mScarlet production. (C) Diagram of the design of P_rrtetO_, a rickettsia-compatible promoter incorporating a tet operator. (D) Fluorescence microscopy images showing the induction of mScarlet fluorescence by SS_L_ pRAMtetS in the presence of aTc. Rickettsia shown in green, mScarlet in magenta, and nuclei in blue. (E) Plasmid map of pRAMin, a generalized inducible expression vector for *R. rickettsii* featuring an NheI site into which a gene of interest can be cloned using NEB HIFI assembly.

### Adaptation of inducible promoter to generate a CRISPRi plasmid

In order to leverage the utility of the aTc inducible promoter system, we sought to generate a single plasmid based CRISPRi system that would be compatible with *R. rickettsii*. We opted to use the d*cas9* derived from *Streptococcus thermophilus* which was shown to be compatible with CRISPRi tools in the context of *Mycobacteria tuberculosis*, borrowing from the design elements described in Rock et al (18). To generate a CRISPRi vector (pRAMci), a *R. rickettsii* codon biased d*cas9_Sth1_* was cloned into the NheI site, with the addition of an adjacent terminator from *rompA* to transcriptionally isolate downstream expression of plasmid elements (Fig 2A, B). Downstream we incorporated a cassette for expression of a chimeric single guide RNA (sgRNA), which combines CRISPR RNA (crRNA) and *trans*-acting RNA (tracrRNA) to target dCas9*_Sth1_* (18, 26). To drive constitutive expression of this cassette, we reused the minimal version of the *rompB* promoter that had been used for the aTc inducible promoter design, but without the tetO operator so that the transcriptional start site could be utilized as the +1 transcription site for sgRNA expression (Fig 2B). Using a rickettsial promoter with a well-defined transcriptional start site may constitute a critical design element because the correct base pairing of the twenty base pairs of the targeting RNA to the predicted target have been shown to be critical for efficient targeting of Cas9 (27). To further ensure the transcriptional termination of the sgRNA, the previously characterized bidirectional terminator from *Rickettsia prowazekii*, P*_RP703-704_* was added downstream (13) (Fig 2B).

**Figure 2:**
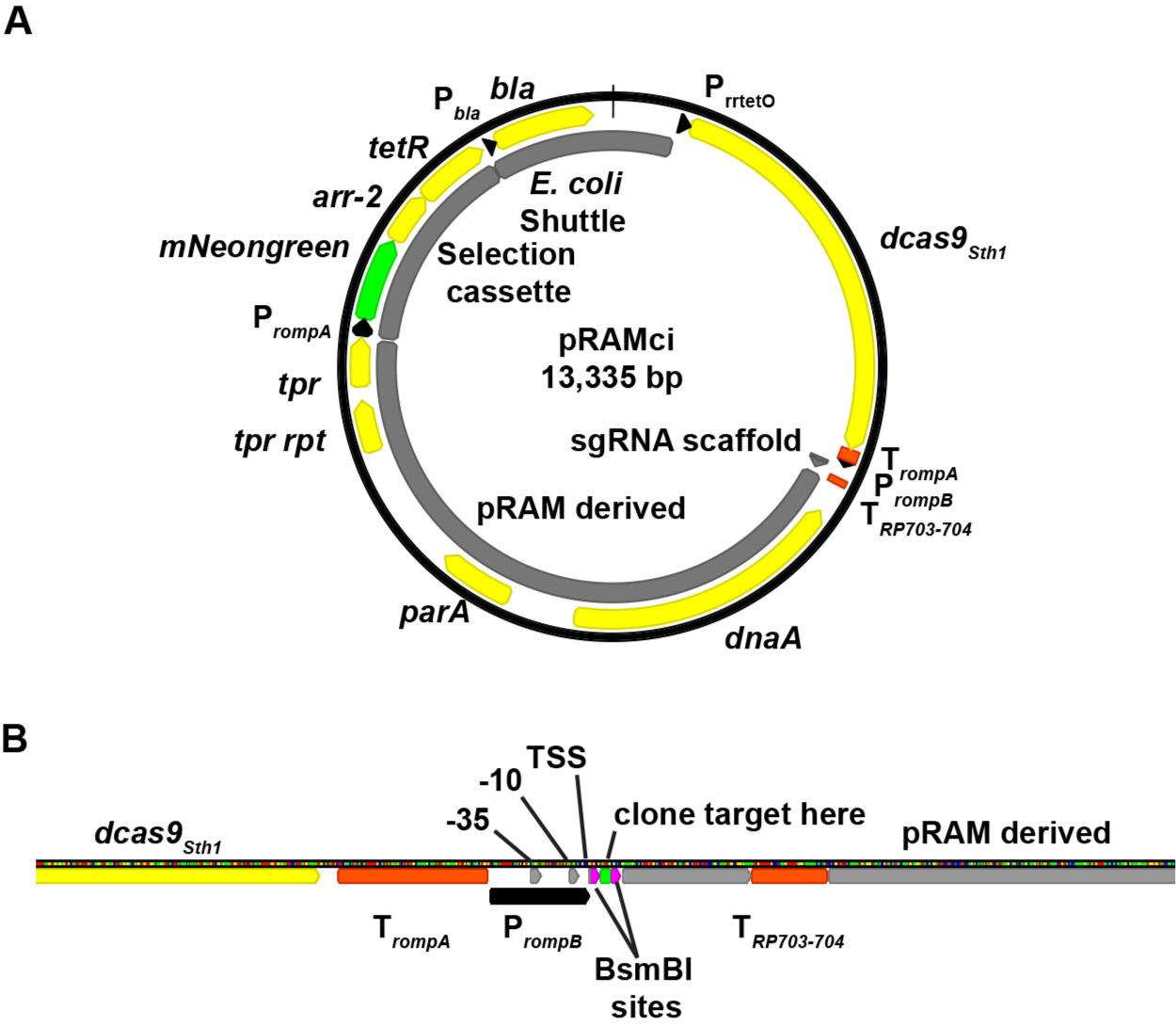
Design of an inducible CRISPRi plasmid targeting *rarP2*. (A) Schematic of pRAMci, a generalized CRISPRi vector for *R. rickettsii*. Key elements include a *R. rickettsii* codon biased d*cas9_Sth1_* cloned into the NheI site and cassette for the targeting of dCas9*_Sth1_*. (B) Diagram of promoter and sgRNA targeting cassette used in pRAMci. Terminators are highlighted in red and annotated as “T” with subscript. Elements of the promoter, P*_rompB_*, include −35, −10, and TSS (transcriptional start site) are shown, as well as the BsmBI sites where 20 bp targeting elements may be cloned in using NEB HIFI assembly.

### Assessment of plasmid platform for inducible expression of d*cas9*_Sth1_ and CRISPR interference capacities

To assess the functionality of the inducible promoter and CRISPRi functionalities of our plasmid system, RT-qPCR was performed to determine gene expression levels when either aTc or DMSO vehicle control was added (Fig 3). *R. rickettsii* strains were used to infect Vero cells at an MOI of 1 for 24 hours before isolation of total RNA, subsequently used to generate cDNA and perform qPCR. Levels of d*cas9_Sth1_* were increased between 11 to 16-fold in all strains harboring a CRISPRi plasmid in response to the addition of 10 ng/mL aTc (Fig 3A). An important note is that in these strains, a basal level of d*cas9_Sth1_* transcript is detected compared to the parental strain, SS_L_, and to negative (no reverse transcriptase) controls in the qPCR experiment. qPCR was also performed on the rickettsial gene *rickA*, previously implicated in early actin-based motility (28–30) (Fig 3B). In each strain expressing d*cas9_Sth1_* under plus aTc, the expression of *rickA* is reduced by approximately 3-fold. This suggests that induction and overexpression of d*cas9_Sth1_* may have a toxic effect, lowering expression of transcripts non-specifically. This observation was borne out by further studies (see below) showing that induction and overexpression of dCas9 displayed off target effects with or without targeting to specific genes. Notably, while SS_L_ and Iowa are not affected by aTc, consistent with plaque recovery data (Fig 1A), strains SS_L_ pRAMci and SS_L_ pRAMci-RARP2 appear elongated when aTc is added which is indicative of stress (Fig S2). This correlates with increased expression of d*cas9_Sth1_* in these strains and a non-specific decrease in tested RNA levels (Fig 3), adding to evidence that overexpression of d*cas9_Sth1_* may have toxic effects. To confirm a possible negative effect of dCas9*_Sth1_* induction on rickettsial growth, a one-step growth curve in the presence or absence of 10 ng/ml aTc was conducted. Induction of dCas9*_Sth1_* expression from either pRAMci (vector control) or pRAMci-RARP2 resulted in approximately a log decrease in *R. rickettsii* replication by 48 hr post-infection. Subsequent studies on expression were limited to the uninduced condition to avoid confounding results potentially arising from dCas9*_Sth1_* overexpression.

**Figure 3:**
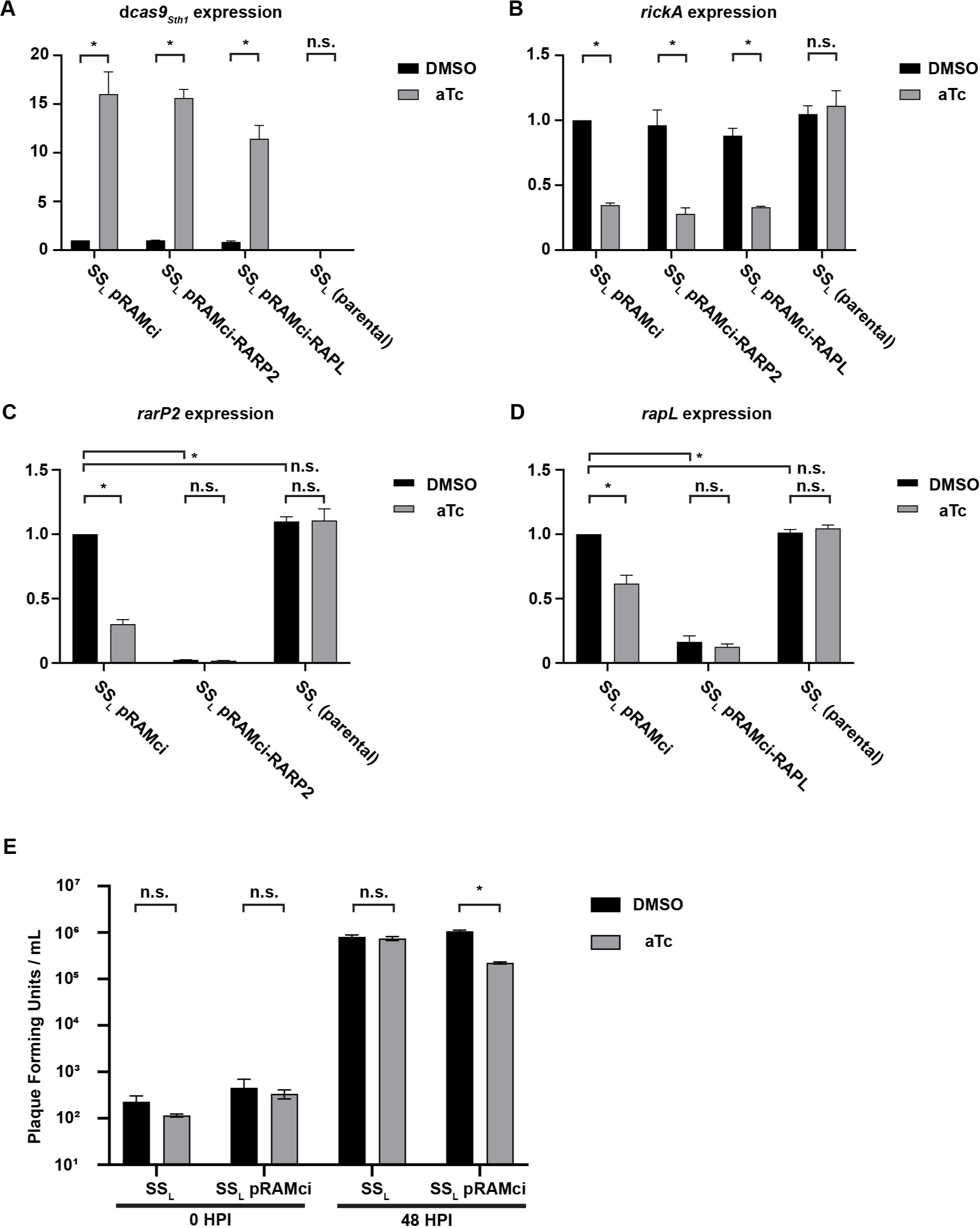
RT-qPCR for RNA expression levels in CRISPRi strains. All panels show expression levels relative to the housekeeping gene, *rpoD*, and normalized to SS_L_ pRAMci in the DMSO control condition. (A) Relative expression levels of d*cas9_Sth1_*. (B) Relative expression levels of *rickA*, a rickettsial gene expected to be unrelated to the genes targeted by the CRISPRi constructs. (C) Relative expression levels of *rarP2*. (D) Relative expression levels of *rapL*. Statistics for RT-qPCR data were performed on fold-expression means derived from three biological replicates which were normalized to SS_L_ pRAMci in the DMSO control condition. Error bars represent standard deviation. perform two-way ANOVA followed by Šídáks multiple comparisons test to compare DMSO vs aTc conditions and followed by Dunnett’s multiple comparisons test to compare the DMSO conditions between strains for C and D. “*” represents and adjusted p value ≤ 0.0001 while “n.s.” (not significant) represents an adjusted p value of ≥ 0.01. (E) Growth of *R. rickettsia* with and without pRAMci in the presence or absence of 10 ng / mL aTc after 48 hours in Vero cells. Data were analyzed via unpaired t test with Welch correction and “*” denotes a P value of 0.001152.

### Inhibition of specific rickettsial genes

To test the efficacy of the CRISPRi system in rickettsia, the rickettsial ankyrin repeat protein 2 (*rarP2*) and rickettsial autotransporter protease lipoprotein (*rapL*) were selected as potential targets due to their known phenotypes and because they are both enzymes which might test the limits of the CRISPRi design (31–33). These genes were scanned for targetable sequences compatible with dCas9*_Sth1_*, which utilizes a protospacer adjacent motif (PAM) sequence of NNAGAAW (34), and contributes to the binding of Cas9 to target DNA (35). To target the sgRNA to *rarP2*, we identified a potential target sequence 5’-AAACCTAAATCAGTAGGTTT-3’ with adjacent PAM site 5’-ACAGAA-3’ near the 5’ end of *rarP2* on the opposing strand (Fig S3). Similarly, to target *rapL* we identified the potential targeting sequence 5’-ATGAGTAGTAGAACACCCTG-3’ with adjacent PAM site 5’-AAAGAAA-3’ also on the opposing strand (Fig S3B).

The expression of *rarP2* in SS_L_ pRAMci-RARP2 (carrying a CRISPRi plasmid targeting *rarP2*) is reduced by more than 97% without the addition of aTc (Fig 3C). Addition of aTc decreased expression slightly but this was not statistically significant different and led to non-specific reduction in rarP2 transcript production in the SS_L_ pRAMci (vector control) and SS_L_ pRAMci-RAPL (CRISPRi plasmid targeting *rapL*) strains. However, the specific and significant reduction of *rarP2* in the SS_L_ pRAMci-RARP2 strain is promising evidence of successful targeting.

Expression of *rapL* by SS_L_ pRAMci-RAPL (carrying a CRISPRi plasmid targeting *rapL*) was reduced by 83% in the absence of induction by aTc (Fig 3D). In the uninduced condition, the only significant transcript reduction in was in rapL expression, suggesting a positive targeting result of the CRISPRi plasmid. Again, non-specific reductions in *rapL* transcript expression were observed when d*cas9_Sth1_* was overexpressed by addition of aTc.

### Disruption of RarP2 function by CRISPRi

RarP2 from the virulent, SS_L_ strain of *R. rickettsii* has previously been shown to localize to the endoplasmic reticulum and specifically disrupt the trans-Golgi network (TGN), while a truncated form from the avirulent Iowa strain does not cause disruption (31, 32). To assess the effectiveness of our CRISPRi platform in the absence of an effective antibody for RarP2, we assayed TGN disruption by immunofluorescence in Vero cells infected with *R. rickettsii* without aTc induction (Fig 4). The parental SS_L_ and SS_L_ pRAMci vector control strains cause disruption of the TGN, while host cells infected with Iowa_op_ contain intact trans-Golgi after 24 hours. Uninduced SS_L_ pRAMci-RARP2 disrupts the TGN with reduced frequency compared to control strains

**Figure 4:**
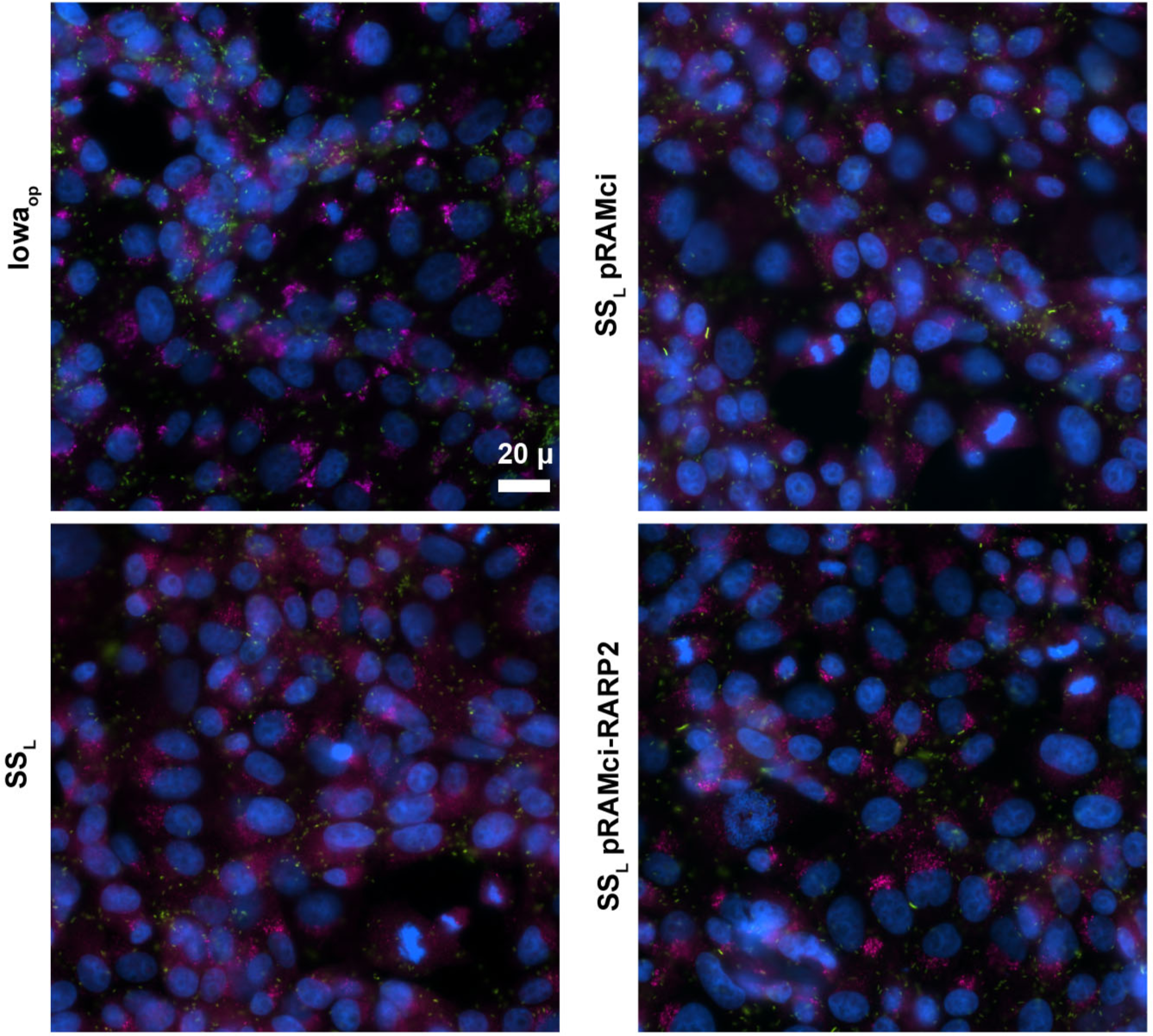
Prevention of trans-Golgi disruption via CRISPRi knockdown of *rarp2* in Vero cells infected with *R. rickettsii*. The *R. rickettsii* Iowa strain expresses a truncated form of *rarP2* which fails to disperse the trans-Golgi, thus serves as a negative control for trans-Golgi dispersion. SS_L_ produces full-length *rarP2* which causes total dispersion of the TGN by 24 hours post infection and serves as a positive control. TGN is shown in red, rickettsiae in green, and nuclei in blue. Bar = 20 μm

When the TGN is disrupted by RarP2, the trans-Golgi marker TGN46 is not properly glycosylated and lower apparent molecular weight forms become apparent by Western blotting (31, 36). Previously, Vero cells infected with Iowa were shown to have TGN46 glycosylation levels identical to uninfected host cells, while infection with SS_L_ caused a glycosylation defect that correlated with TGN dispersion. Here SS_L_ pAN302 has higher levels of glycosylated TGN46 relative to the vector control strain (Fig 5). When treated with deglycosylases, TGN46 in all samples migrate at a lower apparent molecular weight on SDS-PAGE gel (Fig 5). Taken together, these results suggest that the CRISPRi system is capable of disrupting host cell phenotypes precipitated by the rickettsial effector RarP2.

**Figure 5:**
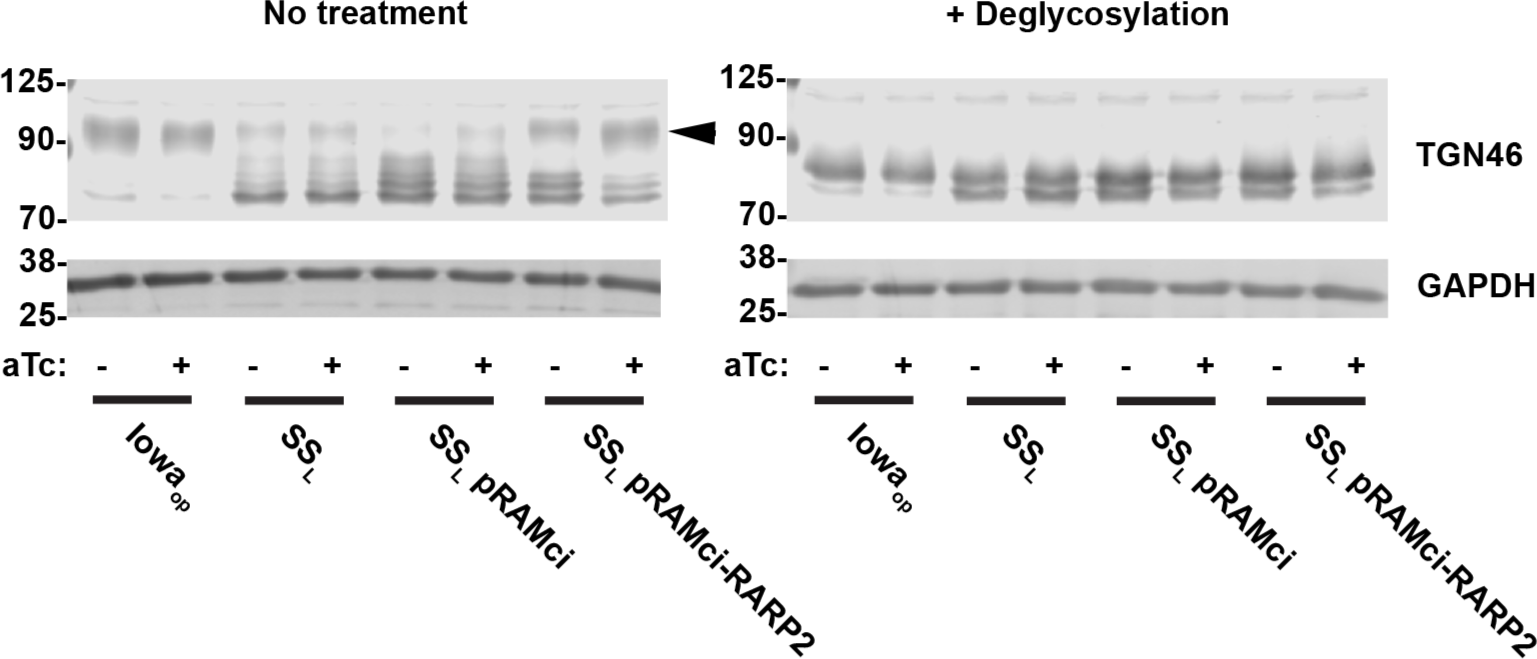
Abrogation of TGN46 deglycosylation by CRISPRi targeting *rarP2*. Samples of Vero cells infected with different rickettsia strains treated with DMSO or aTc were harvested and treated without or with a mixture of deglycosylase enzymes. Western blots for the two sample sets were generated, probing with αTGN46 and αGAPDH as a loading control. Dispersion of the TGN by SS_L_ results in decreased glycosylation of TGN46. The truncated *rarP2* allele expressed by Iowa_op_ fails to cause trans-Golgi dispersion and therefore host cell TGN46 is modified via glycosylation similarly to uninfected cells (31). Arrowhead indicates the major glycosylated form of TGN46.

### Expression of targeting sequence alone is not sufficient to prevent trans-Golgi disruption

To exclude the possibility that an RNAi-like effect is occurring, whereby expression of the sgRNA designed to target *rarP2* might by itself affect transcriptional levels of *rarP2*, Vero cells were infected with *R. rickettsia* strains carrying plasmids that lack d*cas9_Sth1_* (Fig 6). Both SS_L_ pRAMtet (vector control) and SS_L_ pRAMtet-RARP2sgRNA (expressing the sgRNA targeting *rarP2* present in pRAMci-RARP2) failed to prevent disruption of the trans-Golgi by RarP2. This result suggests that dCas9*_Sth1_* is necessary to achieve downregulation of the target genes.

**Figure 6:**
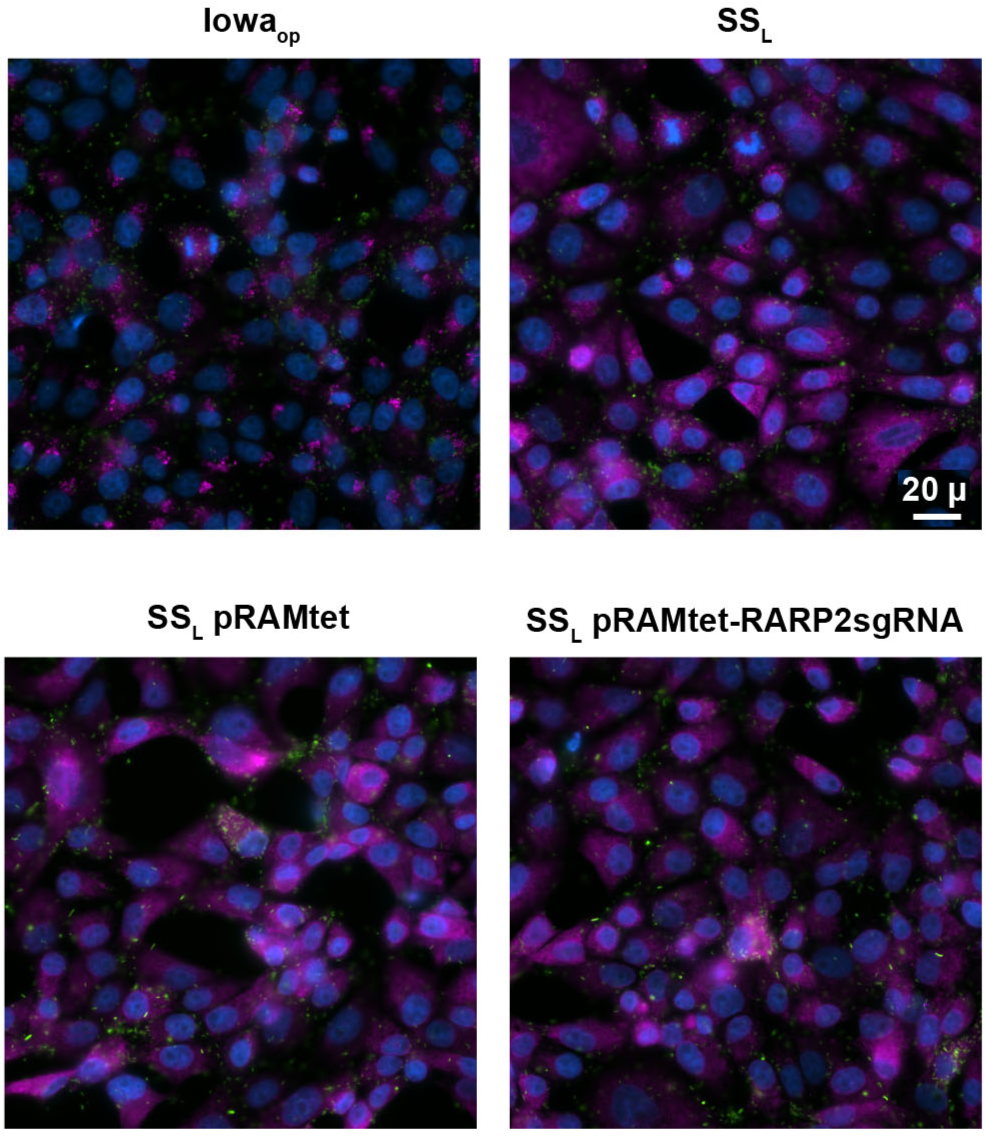
Expression of sgRNA targeting *rarP2* without d*cas9_Sth1_* has no effect on trans-Golgi disruption. Vero cells were infected at 1 MOI for 24 hours at 34°C with *R. rickettsii* carrying plasmids lacking the d*cas9_Sth1_* which is present in pRAMci, with (pRAMtet-sgRNA-RARP2) and without (pRAMtet) the same sgRNA targeting sequence contained in pRAMci-RARP2. The parental SS_L_ strain and Iowa_op_ are included as positive and negative control for trans-Golgi disruption respectively. Rickettsia are visualized in green, the trans-Golgi in magenta, and the nucleus in blue,

### Reduction in RapL expression by CRISPRi

RapL is a putative serine protease lipoprotein that has been implicated in the cleavage of rickettsial autotransporters (33). Although the reduction in *rapL* transcription was less than was observed for the reduction in *rarP2* (Fig 3C, D), we assessed the effects of this reduced expression on the expected protein levels and phenotype. A significant reduction in RapL was observed by Western blot (Fig 7). However, this reduction in expression was not enough to detect uncleaved rOmpB,

**Figure 7:**
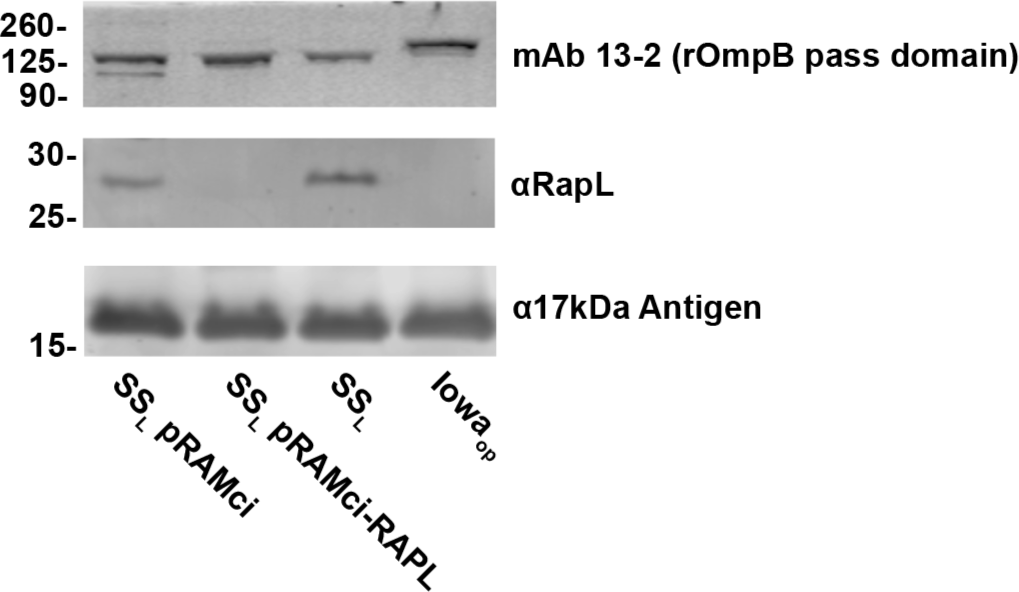
Reduction in RapL by CRISPRi targeting. RapL content is dramatically reduced by CRISPRi but is insufficient to demonstrably reduce rOmpB proteolytic processing.

## DISCUSSION

Largely due to their obligate intracellular nature, genetic tools for *Rickettsia* spp. have lagged behind those available for free-living and facultative intracellular pathogens. Over approximately the past 15 years, tremendous advances have been made. Plasmid expression systems and transposon mutagenesis are now commonly used (2). Despite this progress, directed mutagenesis of specific genes has frequently proven to be difficult. Limited success has been obtained for specific genes under certain conditions, but no robust, widely applicable system has been developed. An LtrA group II intron system (TargeTron) has been used to knock out rOmpA (7) but specific sequences are required in the gene of interest to successfully target it, thus, its use may not be possible. Homologous recombination has been achieved under specific circumstances but selection and isolation of mutants can be difficult (8, 37). Given the somewhat limited genetic tools available for *Rickettsia*, the plasmids described here represent a useful step forward. Inducible expression itself should be a powerful tool for studying the function of the many unknown genes in rickettsia and addressing the occasional issue of toxicity due to overexpression. The simplicity of a single plasmid CRISPRi system should provide facile implementation into any *Rickettsia* species or strain of interest that is normally transformable with pRAM-based vectors, allowing for gene knockdowns to answer a variety of questions. Successful transformation of pRAMdRGA has been shown in variety of rickettsia in addition to *R. rickettsii* including *R. parkeri*, *R. monacensis*, *R. amblyomatis*, *R. montanensis*, *R. bellii*, and *R. typhi* (6, 38, 39).

One caveat to the CRISPRi system presented here is apparent stress to the rickettsia caused by overexpression of d*cas9_Sth1_*. Overexpression of dCas9 can itself be toxic despite being catalytically dead due to proteotoxic effects and off-target binding to essential genes (18, 40). Rock et al (18) approached the problem of proteotoxicity by testing the expression of various dCas9 orthologs for toxicity in *Mycobacteria* before settling on dCas9*_Sth1_*. Other dCas9 orthologs could be tested in rickettsia, and this may have the added benefit of employing different PAM sites which may enable better targeting of certain genes which lack an appropriately placed PAM site compatible with dCas9*_Sth1_*. An elegant strategy used to solve the overexpression issue in the CRISPRi system for *C. trachomatis* was to append dCas9 with a VAA degradation tag, decreasing the overall abundance of dCas9 in the *Chlamydia* (17). If this tag similarly results in increased degradation in rickettsia, it would be an straightforward modification to the current vector.

Another avenue is to address issues with dCas9 overexpression or to generally tune the inducible expression plasmid is to alter the level of expression by the inducible promoter. A perennial issue of the implementation of inducible promoters is the degree to which the promoter is “leaky” in the uninduced state. This has been observed in prokaryotic inducible expression systems including those incorporating *lacI*, *araC*, and *tetR* among others (41–43). One solution may be to simply utilize a lower activity promoter with a similar positioning of tetO, as P*_rompB_* is expected to be highly expressed and thus could contribute to the leakiness of the inducible promoter as designed. Altering the sequence of critical promoter motifs such as the −35, −10, and Shine-Dalgarno sequences is also a common strategy to reduce expression levels of a promoter (44). However, an important note here is that rickettsia may not to utilize a Shine-Dalgarno sequence for transcriptional initiation as most genes were shown not to have a bioinformatically predictable Shine-Dalgarno sequence, despite having a typical anti-Shine-Dalgarno sequence in their 16S ribosomal RNA identical to *E. coli* (45). This may also be accomplished by further transcriptionally isolating the inducible d*cas9*_Sth1_ from the *E. coli* origin by incorporating an additional terminator sequence in between them, which proved critical in the design of CRISPRi plasmids for *Mycobacteria*.

A final concern is one common to all systems of transcriptional downregulation. Protein stability, turnover, and activity are all factors that must be considered. We were able to demonstrate RarP2 interference was sufficient to disrupt the associated phenotype of TGN disruption. However, despite being able to show a decrease in protein levels of RapL by CRISPRi, sufficient enzymatic activity remained that autotransporter processing showed no demonstrable defect. In this case, inactivation or deletion of the gene may be necessary to demonstrate function. We had previously shown that mutation of any of the amino acids of the serine protease catalytic triad in RapL were sufficient to inhibit autotransporter processing (33). Although there is sufficient expression of d*cas9_Sth1_* in this system to cause reduced expression of the target gene in the absence of aTc induction, this might be leveraged as beneficial in animal experiments where it may be difficult to dose with anhydrotetracycline and avoid possible effects from a disturbed microbiome.

This work may be extended by applying other developments in CRISPR technology to further enhance the genetic toolkit available for *Rickettsia*. The adaptation of Cas12a (also referred to as Cpf1) is one possible enhancement (46). Cas12a differs from Cas9 in that Cas12a combines both DNA cleavage activity and the RNase activity used to process CRISPR arrays in one enzyme, which allows the usage of a CRISPR array to silence multiple genes using dCpf1 (47). Inactivating both catalytic sites of Cpf1 would be beneficial as an alternative enzyme to achieve CRISPR interference because its PAM site, TTTV, is more AT-rich than other CRISPR enzymes thus potentially providing more suitable targeting sequence selection for the AT-rich *R. rickettsii* genome. The PAM sites of both Cas9 and Cpf1 have also been mutagenized successfully to increase the diversity of potential targeting sequences for CRISPRi systems (48, 49). CRISPR prime editing (50) utilizes a variant of Cas9 that only nicks a single DNA strand combined with a reverse transcriptase to introduce specific base pair mutations, insertions, and deletions, has recently been applied to *E. coli* (51) and may be possible to apply in *Rickettsia* as well. The ability to knock out expression of non-essential rickettsial genes would prove advantageous.

## MATERIALS AND METHODS

### Propagation of *R. rickettsii* strains and maintenance of Vero cells

*Rickettsia rickettsii* strains Sheila Smith (CP000848.1) and Iowa (CP000766.3), was maintained in Vero cells at 34°C. For purifications, infected cells were lysed by Dounce homogenization, centrifuged through a 30% Renografin pad, washed twice with sucrose, and were stored in brain-heart infusion (BHI) media at −80°C, as previously described (33).

For plaque assays, 10-fold serial dilutions of rickettsia in BHI media, were added to Vero cell monolayers in 6-well plates. Following bacterial invasion, the cells were overlaid with M199 media supplemented with 5% FBS and 0.5% molten low-melt agarose. After 5-7 days of growth at 34°C, plaques were counter stained with thiazolyl blue tetrazolium bromide, 0.3% (MTT). The following day, plaque forming units were counted, and PFU/mL was calculated.

### Transformation of rickettsia

Rickettsia were transformed similarly to previously described (52). Briefly, rickettsia were purified from ten T150 flasks of infected Vero cells as described above. The purified rickettsia were then resuspended in 100 μL BHI, to which 10 μg of plasmid was added. Electroporation was performed using a BioRad Gene Pulser Xcell set at 2.5kV, 200 ohm, and 25uF. Rickettsia were recovered from cuvette in BHI and used to infect 6-well plates with Vero cell monolayers for 30 minutes, then M199 + 2% FBS was added to each well and the plates were incubated at 34°C. After 9 hours of incubation, media was replaced with 3 mL M199 + 5% FBS + 0.5% low-melt agarose + 200 ng/mL rifampin. Plates were incubated at 34°C for 7-9 days when plaques were visible and could be selected and used for propagation.

### Rickettia growth curves and anhydrotetracycline tolerance

To determine tolerance to anhydrotetracycline, SS_L_ was used to infect wells containing confluent Vero cells in 6-well plates overlain with media containing soft agar. SS_L_ was serially diluted in BHI to achieve a dilution of approximately 600 PFU/mL in BHI, of which 100 μL was added per well to infect to achieve approx. 60 plaques per well. After 30 minutes infection at 34°C, wells were overlayed with M199 + 5% FBS mixed with DMSO control or 10, 50, 100, 250, or 500 ng/mL aTc. Plates were incubated for 7 days at 34°C and stained with MTT as previously described () to count the number of plaques.

### Molecular cloning and sequencing

All molecular cloning steps for plasmids were performed using NEB Stable Competent *E. coli* (NEB, C3040). Inserts and fragments generated by PCR were synthesized using Q5 High Fidelity DNA Polymerase (NEB, M0491). All primers were synthesized by Integrated DNA Technologies. Restriction enzymes including BsmBI-v2 (NEB, R0739), NheI-HF (NEB, R3131), and KpnI-HF (NEB, R3142) were used according to the manufacturer’s recommendations. Digested vectors to be used in ligation reactions were treated with recombinant shrimp alkaline phosphatase (NEB, M0371). PCR fragments and digested vectors were purified using GeneJET PCR Cleanup Kit (ThermoFisher, K0701) or GeneJET Gel Extraction Kit (ThermoFisher, K0691). T4 DNA Ligase (NEB, M0202) and NEBuilder DNA HIFI Assembly Master Mix (NEB, 2621) were used to combine vectors and inserts. Plasmids were purified using GeneJET Plasmid Miniprep Kit (ThermoFisher, K0502) and GeneJET (ThermoFisher, K0492), and quantified using a Nanodrop 2000 Spectrophotometer (ThermoFisher) and or Qubit Flex Fluorometer (ThermoFisher) with Qubit dsDNA Quantification Assay Kit (ThermoFisher, Q32850). Sanger sequencing on initial cloning steps was performed by AGCT Inc. Whole plasmid sequencing using Oxford Nanopore technology with custom analysis and annotation was performed by Plasmidsaurus.

### Construction of an inducible expression system

Modifications to the original pRAM18dRGA[MCS] vector were needed to achieve downstream goals (Fig S1). The high copy *E. coli* origin of replication in pRAM18dRGA[MCS] was replaced with a lower copy number *E. coli* origin of replication by amplifying the pMB1 origin and *bla* from pMAL-C2x and using splice overlap extension PCR to combine these pieces to exclude the unneeded M13 phage origin and generate pRAM18lc. This simultaneously removed two BsmBI restriction sites which allowed for BsmBI sites to be incorporated into later cloning steps. The change of origins reduces the amount of plasmid purified from *E. coli* via kit by about half. pRAM18lc was further reduced in size by redesigning the rifampin resistance and fluorescent protein cassette for selection in rickettsia by removing the *rpsL* promoter and combining *R. rickettsii* codon optimized mNeongreen into a polycistron with the original *R. prowazekii* codon optimized rifampin resistance gene, *arr-2*, driven by the *rompA* promoter to generate pRAM18lcN2.

To test and build an inducible expression vector, an intermediate construct was generated by incorporating *R. rickettsii* codon optimized *tetR* to the end of the polycistron. Codon optimized mScarlet was inserted into the multiple cloning site, such that different promoters could be tested by HIFI assembly at an NheI site adjacent to the start codon of mScarlet. pRAMtetS was generated by cloning the designed P_rrtetO_ described in Figure 1 to allow detection of mScarlet fluorescence in response to anhydrotetracycline. To generate a generalized inducible expression vector, a sequence containing P_rrtetO_, an NheI site for HIFI cloning, and a predicted terminator from *rompA* was synthesized and cloned into pRAMtet to create pRAMin.

### Construction of CRISPRi plasmids

To generate CRISPRi vectors, a *R. rickettsii* codon optimized d*cas9_Sth1_* was cloned into the NheI site of pRAMin. A generalized sgRNA sequence for dCas9*_Sth1_*, including BsmBI sites to clone targeting sequences was combined with a minimal promoter from *rompB* to drive expression. An RP703-704 terminator sequence was designed and synthesized by Genscript Inc and cloned into the KpnI site via HIFI assembly and to generate pRAMci. Targeting sequences for *rapL* and *rarP2* were ordered as annealed oligos with HIFI compatible overhangs from Integrated DNA Technologies and combined with BsmBI digested pRAMci to generate plasmids for targeted knockdown.

### Isolation of RNA from *R. rickettsii* infected Vero cells and RT-qPCR

Confluent Vero cells were grown in 24-well plates and infected with of rickettsia at an MOI of 1. One hr after infection, media in each well was replaced with M199 + 2 % FBS + DMSO or 10 ng/mL aTc in DMSO. Plates were incubated for 24 hours at 34°C. Media was removed and of Trizol LS (ThermoFisher Scientific, 10296010) was added to each well and incubated at room temperature for 10 minutes before transferring to microcentrifuge tubes. Samples were processed using the Direct-zol Miniprep Plus Kit (Zymo Research, R2070) with optional DNase I step. Samples were treated with DNase I (New England Biolabs, M0303) and applied to a RNA Clean and Concentrate-5 Kit (Zymo Research, R1013) with the optional DNase I treatment. RNA was quantified on a Nanodrop 2000 (ThermoFisher Scientific) and 200 ng of RNA from each sample was used to generate cDNA using Superscript IV First-Strand Synthesis System (ThermoFisher Scientific, 18091050) with random hexemers. An RNase H step was added and the resulting cDNA sample was diluted with molecular grade H_2_O. Quantitative PCR was performed on samples by assembling the individual reactions in MicroAmp Fast Optical 96-well Reaction Plates (Applied Biosystems, 4346906) as follows: 10 μL 2x SYBR Green PCR Master Mix (Applied Biosystems, 4309155), 0.25 μL each of two relevant 20 μM primers, 7.5 μL of molecular grade H_2_O, and 2 μL cDNA. Plates were sealed using MicroAmp Optical Adhesive Film (Applied Biosystems, 4311971) and briefly centrifuged for 1 minute at 1000 rpm using a M-20 rotor (Thermo Scientific, 75003624) in a tabletop centrifuge (Legend X1R, Sorvall). The experiment was performed on a StepOnePlus Real-Time PCR system (Applied Biosystems) using the following program: pre-incubation 95°C for 10 minutes, an amplification cycle of 95°C for 15 seconds, 56°C for 30 seconds, and extension at 60°C for 30 seconds repeated for 40 cycles, followed by a continuous melting curve. Each individual run included rickettsial genomic DNA (*R. rickettsii* Sheila Smith prepared as previously reported using DNeasy Blood and Tissue Kit (Qiagen, 69504) or relevant plasmid at 1 ng/μL as a positive control for each primer set, water negative controls, and no reverse transcriptase negative controls. Data was analyzed with StepOne Software v2.3 (Applied Biosystems) and GraphPad Prism 9 (Dotmatics). Primers used for RT-qPCR were validated both via melt curve and running PCR products on agarose gel to ensure a single product of the predicted size was generated.

### Immunofluorescence assay and microscopy

Confluent Vero cells were grown in 24-well plates with 12 mm circular #1.5 coverslips (Karl Hecht, 92100105019) and infected with rickettsia at an MOI of 1. Cultures were incubated for 24 hours at 34°C, and inactivated by removing media from wells, rinsing wells in 500 μL HBSS (Gibco, 14025092), then adding 4% paraformaldehyde (Ted Pella, 18505) in PBS for 20 minutes at room temperature. Cultures were rinsed once with 500 μL PBS and permeabilized with 0.1% Triton X-100 in PBS l for 15 minutes at room temperature. Cultures were blocked with 1% bovine serum albumin in PBS for 1 hour at room temperature. Coverslips were labeled with 1 μg/mL mAb 13-2 (rOmpB) (53) and 1 μg/mL rabbit polyclonal αTGN46 (Invitrogen, PA5-23068) in PBS overnight at 4°C. Coverslips were washed three times with PBS. Secondary antibodies included Goat Anti-Mouse IgG H&L (Alexa Fluor® 488) (Abcam, ab150117) and Goat Anti-Rabbit IgG H&L (Alexa Fluor® 568) (Abcam, ab175695) diluted 1:550 in PBS were added and incubated at room temperature for three hours. Supernatent was removed from wells and 1 μM Hoechst 33342 (Thermo Scientific, 62249) in PBS was added and incubated for 10 minutes at room temperature. Coverslips were washed four times with PBS, then mounted on slides using ProLong Diamond antifade mounting fluid (Invitrogen, P36961). Specimens were visualized using a Nikon Eclipse Ti2 set up for widefield microscopy with a Nikon DS-Qi2 16.25MP sCMOS Camera, DAPI Filter Set (Nikon, 96370), GFP/FITC/Cy2 Filter Set (Nikon, 96372), DSRed/TRITC/Cy3 Filter Set (Nikon, 96374), CFI60 Plan Apochromat Lambda 60x Oil Immersion Objective Lens, N.A. 1.4, W.D.0.13mm, F.O.V. 25mm, DIC (Nikon, MRD01605) and Lumencor SOLA LED Epifluorescence Light Source (Nikon, 77060113). Three biological replicates of each sample were imaged at five randomized points each using autofocus with the DAPI filter set to acquire representative images using Nikon Elements AR version 5.11.02 software.

### Deglycosylation assay and Western blot

Deglycosylation assays were performed as described previously (31). Confluent Vero cells in a 24-well plate were infected with rickettsia at an MOI of 1 in M199-2 media containing DMSO as a vehicle control, or 10 ng/mL aTc for 24 hours at 34°C. Media was aspirated and wells were washed with molecular grade water and scraped into 150 μL molecular grade water. Samples were shaken vigorously for 10 minutes using a tabletop vortex mixer, then 40 μL of each sample was used reactions as outlined in the manufacturer’s protocol for denaturing reaction conditions using the Protein Deglycosylation Mix II Kit (New England Biolabs, P6044) both with and without deglycosylase enzymes. After 1 hour incubation at 37°C, an equal volume of 2x Laemmli sample buffer was added to each reaction and samples were incubated at 99°C for 10 minutes and loaded onto an 8% acrylamide SDS-PAGE gel and run at 80 volts. Protein was transferred to Immobilon-FL PVDF membrane (Merck Millipore, IPFL00010) at 100 volts using 1x Towbin transfer buffer with 20% methanol. After drying overnight, the membrane was rehydrated in methanol, briefly rinsed in water, washed in TBS buffer, and blocked for 1 hour at room temperature in Intercept PBS blocking buffer (Li-cor, 927-70001). Primary antibodies αTGN46 (Invitrogen, PA5-23068) and αGAPDH (Abcam, ab83956) were diluted to 1 μg/mL each in Intercept PBS blocking buffer + 0.2% tween 20 and incubated for 3 hours at room temperature. The membrane was washed four times in TBS-T buffer. Secondary antibodies IRDye 800CW Donkey anti-chicken (Li-cor, 926-32218) and IRDye 680RD Goat anti-Rabbit (Li-cor, 926-68071) were diluted 1:10,000 in Intercept PBS blocking buffer + 0.2% tween 20 + 0.02% SDS and incubated with membrane for 1 hour at room temperature. The meembrane was washed four more times in TBS-T, and once in TBS before visualization on an Odyssey Clx Imager (Li-cor).

### Plasmid sequences

Plasmid sequences have been deposited in GenBank under accession numbers: XXXXX, XXXXX, XXXXX, etc.

### Statistics

GraphPad Prism 9.3.1 was used for all statistical analyses.

## ACKNOWLEDGEMENTS

This work was supported by the Division of Intramural Research, NIAID/. We thank Uli Munderloh and her laboratory for providing the pRAM18dRGA[MCS] shuttle vector.

## Supplemental Figure Legends

**Figure S1:**
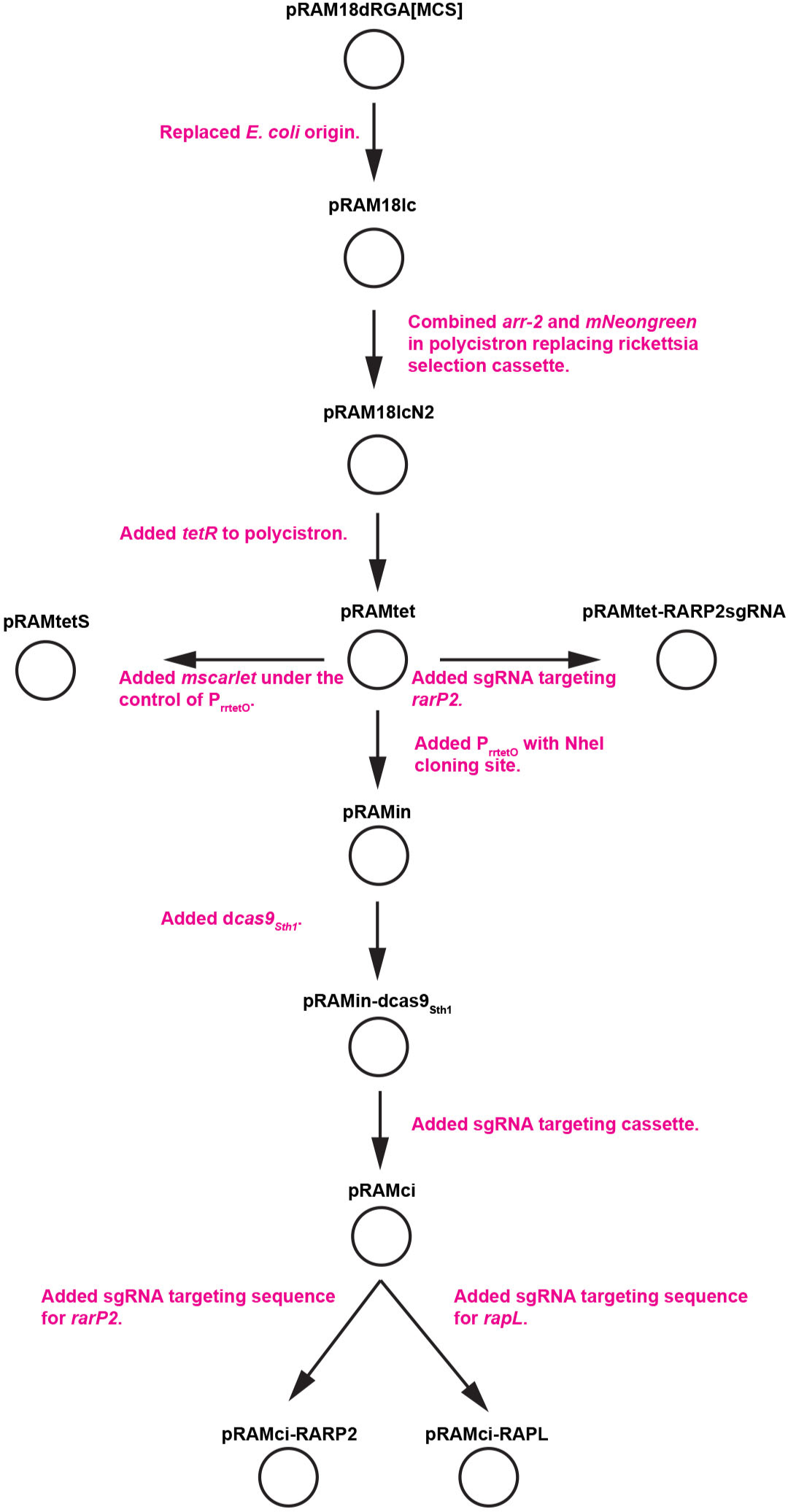
Cloning scheme for critical plasmids used in this study.

**Figure S2:**
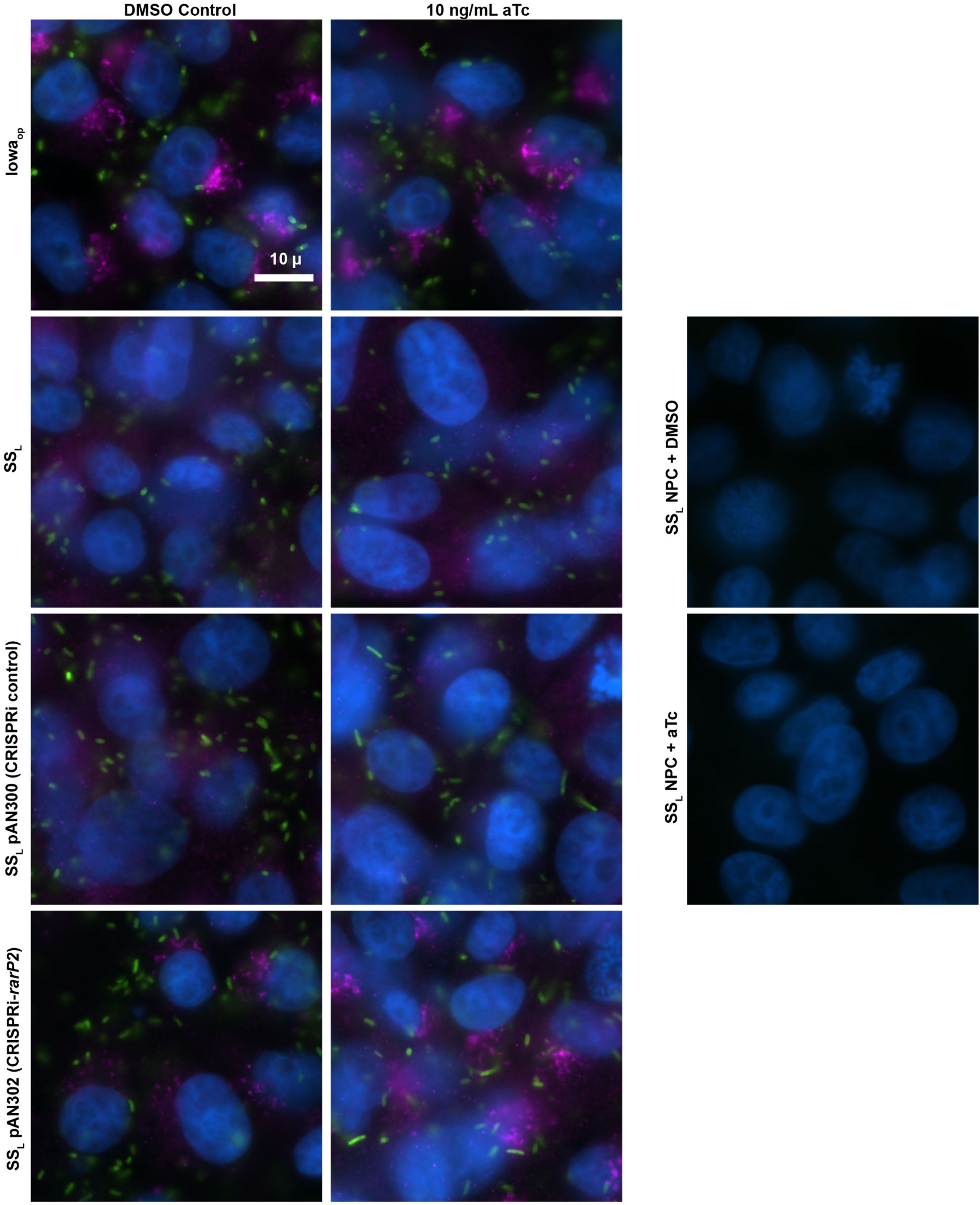
Schematic of *rarP2* and *rapL* with sgRNA targeting sequence (magenta) and PAM sequence (blue) labeled.

**Figure S3:**
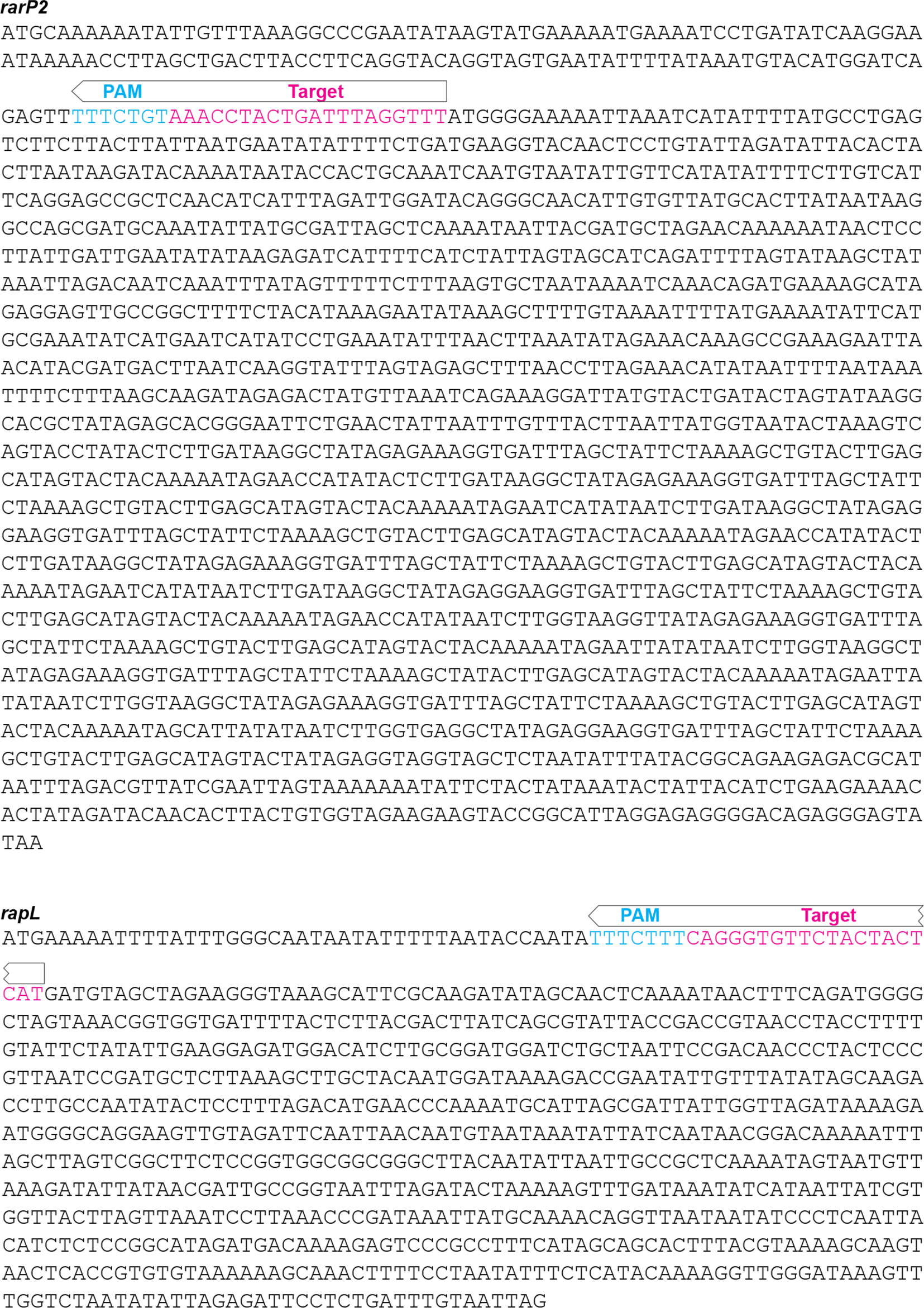
Elongation of rickettsia due to dc*as9_Sth1_* expression. Experiment is identical to Fig 4 with induction of dCas9 by 10 ng/ml aTc, shown at higher magnification so that elongated rickettsia are evident.

## Notes

### Competing Interest Statement

The authors have declared no competing interest.

## REFERENCES

1. McClure EE, Chavez ASO, Shaw DK, Carlyon JA, Ganta RR, Noh SM, Wood DO, Bavoil PM, Brayton KA, Martinez JJ, McBride JW, Valdivia RH, Munderloh UG, Pedra JHF. 2017. Engineering of obligate intracellular bacteria: progress, challenges and paradigms. Nat Rev Microbiol 15:544–558.

2. Fisher DJ, Beare PA. 2023. Recent advances in genetic systems in obligate intracellular human-pathogenic bacteria. Front Cell Infect Microbiol 13:1202245.

3. Rachek LI, Tucker AM, Winkler HH, Wood DO. 1998. Transformation of Rickettsia prowazekii to rifampin resistance. J Bacteriol 180:2118–24.

4. Liu ZM, Tucker AM, Driskell LO, Wood DO. 2007. Mariner-based transposon mutagenesis of Rickettsia prowazekii. Appl Environ Microbiol 73:6644–9.

5. Baldridge GD, Burkhardt NY, Felsheim RF, Kurtti TJ, Munderloh UG. 2007. Transposon insertion reveals pRM, a plasmid of Rickettsia monacensis. Appl Environ Microbiol 73:4984–95.

6. Burkhardt NY, Baldridge GD, Williamson PC, Billingsley PM, Heu CC, Felsheim RF, Kurtti TJ, Munderloh UG. 2011. Development of shuttle vectors for transformation of diverse Rickettsia species. PLoS One 6:e29511.

7. Noriea NF, Clark TR, Hackstadt T. 2015. Targeted knockout of the Rickettsia rickettsii OmpA surface antigen does not diminish virulence in a mammalian model system. MBio 6.

8. Nock AM, Clark TR, Hackstadt T. 2022. Regulator of actin-based motility (RoaM) downregulates actin tail formation by *Rickettsia rickettsii* and is negatively selected in mammalian cell culture. mBio 13:e0035322.

9. Oliver JD, Burkhardt NY, Felsheim RF, Kurtti TJ, Munderloh UG. 2014. Motility characteristics are altered for Rickettsia bellii transformed to overexpress a heterologous rickA gene. Appl Environ Microbiol 80:1170–6.

10. Cai J, Pang H, Wood DO, Winkler HH. 1995. The citrate synthase-encoding gene of Rickettsia prowazekii is controlled by two promoters. Gene 163:115–9.

11. Policastro PF, Hackstadt T. 1994. Differential activity of *Rickettsia rickettsii ompA* and *ompB* promoter regions in a heterologous reporter gene system. Microbiol 140:2941–2949.

12. Qin A, Tucker AM, Hines A, Wood DO. 2004. Transposon mutagenesis of the obligate intracellular pathogen Rickettsia prowazekii. Appl Environ Microbiol 70:2816–22.

13. Woodard A, Wood DO. 2011. Analysis of convergent gene transcripts in the obligate intracellular bacterium Rickettsia prowazekii. PLoS One 6:e16537.

14. de la Torre JC, Ortin J, Domingo E, Delamarter J, Allet B, Davies J, Bertrand KP, Wray LV, Jr., Reznikoff WS. 1984. Plasmid vectors based on Tn10 DNA: gene expression regulated by tetracycline. Plasmid 12:103–10.

15. Qi LS, Larson MH, Gilbert LA, Doudna JA, Weissman JS, Arkin AP, Lim WA. 2013. Repurposing CRISPR as an RNA-guided platform for sequence-specific control of gene expression. Cell 152:1173–83.

16. Wachter S, Cockrell DC, Miller HE, Virtaneva K, Kanakabandi K, Darwitz B, Heinzen RA, Beare PA. 2022. The endogenous Coxiella burnetii plasmid encodes a functional toxin-antitoxin system. Mol Microbiol 118:744–764.

17. Ouellette SP, Blay EA, Hatch ND, Fisher-Marvin LA. 2021. CRISPR Interference To Inducibly Repress Gene Expression in Chlamydia trachomatis. Infect Immun 89:e0010821.

18. Rock JM, Hopkins FF, Chavez A, Diallo M, Chase MR, Gerrick ER, Pritchard JR, Church GM, Rubin EJ, Sassetti CM, Schnappinger D, Fortune SM. 2017. Programmable transcriptional repression in mycobacteria using an orthogonal CRISPR interference platform. Nat Microbiol 2:16274.

19. Bauler LD, Hackstadt T. 2014. Expression and targeting of secreted proteins from Chlamydia trachomatis. J Bacteriol 196:1325–34.

20. LoVullo ED, Miller CN, Pavelka MS, Jr., Kawula TH. 2012. TetR-based gene regulation systems for Francisella tularensis. Appl Environ Microbiol 78:6883–9.

21. Wickstrum J, Sammons LR, Restivo KN, Hefty PS. 2013. Conditional Gene Expression in Chlamydia trachomatis Using the Tet System. PLoS ONE 8:e76743.

22. Shaner NC, Lambert GG, Chammas A, Ni Y, Cranfill PJ, Baird MA, Sell BR, Allen JR, Day RN, Israelsson M, Davidson MW, Wang J. 2013. A bright monomeric green fluorescent protein derived from Branchiostoma lanceolatum. Nat Methods 10:407–9.

23. Bindels DS, Haarbosch L, van Weeren L, Postma M, Wiese KE, Mastop M, Aumonier S, Gotthard G, Royant A, Hink MA, Gadella TW, Jr. 2017. mScarlet: a bright monomeric red fluorescent protein for cellular imaging. Nat Methods 14:53–56.

24. Whetstine CR, Slusser JG, Zuckert WR. 2009. Development of a single-plasmid-based regulatable gene expression system for Borrelia burgdorferi. Appl Environ Microbiol 75:6553–8.

25. Gibson DG, Young L, Chuang RY, Venter JC, Hutchison CA, 3rd, Smith HO. 2009. Enzymatic assembly of DNA molecules up to several hundred kilobases. Nat Methods 6:343–5.

26. Jinek M, Chylinski K, Fonfara I, Hauer M, Doudna JA, Charpentier E. 2012. A programmable dual-RNA-guided DNA endonuclease in adaptive bacterial immunity. Science 337:816–21.

27. Zhang D, Zhang H, Li T, Chen K, Qiu JL, Gao C. 2017. Perfectly matched 20-nucleotide guide RNA sequences enable robust genome editing using high-fidelity SpCas9 nucleases. Genome Biol 18:191.

28. Gouin E, Egile C, Dehoux P, Villiers V, Adams J, Gertler F, Li R, Cossart P. 2004. The RickA protein of *Rickettsia conorii* activates the Arp2/3 complex. Nature 427:457–461.

29. Jeng RL, Goley ED, D’Alessio JA, Chaga OY, Svitkina TM, Borisy GG, Heinzen RA, Welch MD. 2004. A Rickettsia WASP-like protein activates the Arp2/3 complex and mediates actin-based motility. Cell Microbiol 6:761–769.

30. Reed SCO, Lamason RL, Risca VI, Abernathy E, Welch MD. 2014. Rickettsia actin-based motility occurs in distinct phases mediated by different actin nucleators. Curr Biol 24:98–103.

31. Aistleitner K, Clark T, Dooley C, Hackstadt T. 2020. Selective fragmentation of the trans-Golgi apparatus by Rickettsia rickettsii. PLoS Pathog 16:e1008582.

32. Lehman SS, Noriea NF, Aistleitner K, Clark TR, Dooley CA, Nair V, Kaur SJ, Rahman MS, Gillespie JJ, Azad AF, Hackstadt T. 2018. The Rickettsial Ankyrin Repeat Protein 2 Is a Type IV Secreted Effector That Associates with the Endoplasmic Reticulum. MBio 9.

33. Nock AM, Aistleitner K, Clark TR, Sturdevant D, Ricklefs S, Virtaneva K, Zhang Y, Gulzar N, Redekar N, Roy A, Hackstadt T. 2023. Identification of an autotransporter peptidase of Rickettsia rickettsii responsible for maturation of surface exposed autotransporters. PLoS Pathog 19:e1011527.

34. Horvath P, Romero DA, Coute-Monvoisin AC, Richards M, Deveau H, Moineau S, Boyaval P, Fremaux C, Barrangou R. 2008. Diversity, activity, and evolution of CRISPR loci in Streptococcus thermophilus. J Bacteriol 190:1401–12.

35. Anders C, Niewoehner O, Duerst A, Jinek M. 2014. Structural basis of PAM-dependent target DNA recognition by the Cas9 endonuclease. Nature 513:569–73.

36. Prescott AR, Lucocq JM, James J, Lister JM, Ponnambalam S. 1997. Distinct compartmentalization of TGN46 and beta 1,4-galactosyltransferase in HeLa cells. Eur J Cell Biol 72:238–46.

37. Driskell LO, Yu XJ, Zhang L, Liu Y, Popov VL, Walker DH, Tucker AM, Wood DO. 2009. Directed mutagenesis of the Rickettsia prowazekii pld gene encoding phospholipase D. Infect Immun 77:3244–8.

38. Burkhardt NY, Price LD, Wang XR, Heu CC, Baldridge GD, Munderloh UG, Kurtti TJ. 2022. Examination of Rickettsial Host Range for Shuttle Vectors Based on dnaA and parA Genes from the pRM Plasmid of Rickettsia monacensis. Appl Environ Microbiol 88:e0021022.

39. Hauptmann M, Burkhardt N, Munderloh U, Kuehl S, Richardt U, Krasemann S, Hartmann K, Krech T, Fleischer B, Keller C, Osterloh A. 2017. GFPuv-Expressing Recombinant Rickettsia typhi: a Useful Tool for the Study of Pathogenesis and CD8(+) T Cell Immunology in R. typhi Infection. Infect Immun 85.

40. Rostain W, Grebert T, Vyhovskyi D, Pizarro PT, Tshinsele-Van Bellingen G, Cui L, Bikard D. 2023. Cas9 off-target binding to the promoter of bacterial genes leads to silencing and toxicity. Nucleic Acids Res 51:3485–3496.

41. Muller J, Oehler S, Muller-Hill B. 1996. Repression of lac promoter as a function of distance, phase and quality of an auxiliary lac operator. J Mol Biol 257:21–9.

42. Meisner J, Goldberg JB. 2016. The Escherichia coli rhaSR-PrhaBAD Inducible Promoter System Allows Tightly Controlled Gene Expression over a Wide Range in Pseudomonas aeruginosa. Appl Environ Microbiol 82:6715–6727.

43. Bertram R, Hillen W. 2008. The application of Tet repressor in prokaryotic gene regulation and expression. Microb Biotechnol 1:2–16.

44. Chen Y, Ho JML, Shis DL, Gupta C, Long J, Wagner DS, Ott W, Josic K, Bennett MR. 2018. Tuning the dynamic range of bacterial promoters regulated by ligand-inducible transcription factors. Nat Commun 9:64.

45. Ma J, Campbell A, Karlin S. 2002. Correlations between Shine-Dalgarno sequences and gene features such as predicted expression levels and operon structures. J Bacteriol 184:5733–45.

46. Zetsche B, Gootenberg JS, Abudayyeh OO, Slaymaker IM, Makarova KS, Essletzbichler P, Volz SE, Joung J, van der Oost J, Regev A, Koonin EV, Zhang F. 2015. Cpf1 is a single RNA-guided endonuclease of a class 2 CRISPR-Cas system. Cell 163:759–71.

47. Li M, Chen J, Wang Y, Liu J, Huang J, Chen N, Zheng P, Sun J. 2020. Efficient Multiplex Gene Repression by CRISPR-dCpf1 in Corynebacterium glutamicum. Front Bioeng Biotechnol 8:357.

48. Kleinstiver BP, Prew MS, Tsai SQ, Topkar VV, Nguyen NT, Zheng Z, Gonzales AP, Li Z, Peterson RT, Yeh JR, Aryee MJ, Joung JK. 2015. Engineered CRISPR-Cas9 nucleases with altered PAM specificities. Nature 523:481–5.

49. Gao L, Cox DBT, Yan WX, Manteiga JC, Schneider MW, Yamano T, Nishimasu H, Nureki O, Crosetto N, Zhang F. 2017. Engineered Cpf1 variants with altered PAM specificities. Nat Biotechnol 35:789–792.

50. Anzalone AV, Randolph PB, Davis JR, Sousa AA, Koblan LW, Levy JM, Chen PJ, Wilson C, Newby GA, Raguram A, Liu DR. 2019. Search-and-replace genome editing without double-strand breaks or donor DNA. Nature 576:149–157.

51. Tong Y, Jorgensen TS, Whitford CM, Weber T, Lee SY. 2021. A versatile genetic engineering toolkit for E. coli based on CRISPR-prime editing. Nat Commun 12:5206.

52. Kleba B, Clark TR, Lutter EI, Ellison DW, Hackstadt T. 2010. Disruption of the *Rickettsia rickettsii* Sca2 autotransporter inhibits actin-based motility. Infect Immun 78:2240–7.

53. Anacker RL, Mann RE, Gonzales C. 1987. Reactivity of monoclonal antibodies to *Rickettsia rickettsii* with spotted fever and typhus group rickettsiae. J Clin Microbiol 25:167–71.

